# High-resolution mapping of forest vulnerability to wind for disturbance-aware forestry and climate change adaptation

**DOI:** 10.1101/666305

**Authors:** Susanne Suvanto, Mikko Peltoniemi, Sakari Tuominen, Mikael Strandström, Aleksi Lehtonen

## Abstract

Windstorms cause major disturbances in European forests and forest management can play a key role in making forests more persistent to disturbances. However, better information is needed to support decision making that effectively accounts for wind disturbances. Here we show how empirical probability models of wind damage, combined with existing spatial datasets, can be used to provide fine-scale spatial information about disturbance probability over large areas. First, we created stand-level damage probability models with predictors describing forest characteristics, recent forest management history and local wind, soil, site and climate conditions. We tested three different methods for creating the damage probability models - generalized linear models (GLM), generalized additive models (GAM) and boosted regression trees (BRT). Then, the damage probability maps were calculated by combining the models (GLM, GAM and BRT) with GIS data sets representing the model predictors. Finally, we demonstrated the predictive performance of the maps with a large, independent test data, which shows that the damage probability maps are able to identify vulnerable forests also in new wind damage events (AUC > 0.7). Use of the more complex methods (GAM and BRT) was not found to improve the predictive performance of the map compared to GLM, and therefore we would suggest using the more simple GLM method that can be more easily interpreted. The map allows identification of vulnerable forest areas in high spatial resolution (16 × 16 m^2^ raster resolution), making it useful in assessing the vulnerability of individual forest stands when making management decisions. The map is also a powerful tool for communicating disturbance risks to forest owners and managers and it has the potential to steer forest management practices to a more disturbance aware direction. Our study showed that in spite of the inherent stochasticity of the wind and damage phenomena at all spatial scales, it can be modelled with good accuracy across large spatial scales when existing ground and earth observation data sources are combined smartly. With improving data quality and availability, map-based risk assessments can be extended to other regions and other disturbance types.

## 1. Introduction

Forest wind disturbances have major economic, societal and ecological consequences in Europe. Forest disturbances have substantial effects on forest productivity and carbon storage (Reyer et al., 2017; Seidl et al., 2014), and therefore actions to reduce and manage the disturbances are crucial in assuring the persistence of the forest carbon sinks. The damage caused by wind storms in European forests has increased during the past century (Gregow et al., 2017; Schelhaas et al., 2003; Seidl et al., 2011) and this trend is expected to continue (Ikonen et al., 2017; Seidl et al., 2017). The question of forest wind disturbances is therefore becoming increasingly important in the future.

Forest management practices play a key role in making forests less vulnerable to wind disturbances. Management driven changes in European forests, such as increasing standing timber volume and promotion of conifer species, have been identified as one of the major causes of increased forest disturbances in Europe during the latter half of the 20th century (Schelhaas et al., 2003; Seidl et al., 2011). If management practices are shifted to reduce forest vulnerability to wind, it may be possible to decrease the negative effects of wind disturbances. However, changing the forest management practises to more disturbance-aware direction is not always easy, as illustrated by the 2005 storm Gudrun in southern Sweden. Despite the massive damage and economic losses caused by the storm and the Swedish Forest Agency’s recommendations for alternative, less vulnerable, management options, the forest management practises in the area remained largely unchanged after the storm (Andersson et al., 2018; Valinger et al., 2014). This demonstrates that not only is information about the wind damage risks urgently needed to account for disturbances in management decisions, but it is also crucial that this information is in a form that can be effectively used and communicated to forest owners and managers.

The development of remote sensing methods and the progress of open data policies have substantially increased the amount, quality and availability of spatial data relating to forests. This opens new possibilities for detailed spatial estimation of forest sensitivity to disturbances. Vulnerability of forests to wind damage is affected by forest characteristics, forest management as well as the abiotic environment, such as local wind and soil conditions (Mitchell, 2013). For example, probability of wind damage has been shown to increase with tree height and certain species, such as Norway spruce, are particularly vulnerable to wind (Dobbertin, 2002; Peltola et al., 1999; Valinger and Fridman, 2011). Forest management has major effects on wind damage sensitivity, as trees that have grown in sheltered conditions and have later been exposed to wind, because of thinning or clear cut of the neighboring stand, are especially sensitive to damage (Lohmander and Helles, 1987; Peltola et al., 1999; Suvanto et al., 2016). Areas that are exposed to strong wind gusts (Schindler et al., 2016) or where rooting conditions are limited due to soil characteristics (Nicoll et al., 2006) are more predisposed to wind damage. Therefore, in order to provide useful information on forest vulnerability to wind damage, information from several different sources, scales and disciplines needs to be brought together.

Logistic generalized linear models (GLM) have long been applied in statistical modelling of forest wind damage (Lohmander and Helles, 1987; Suvanto et al., 2016; Valinger and Fridman, 1997). In addition, different approaches allowing more flexible model behaviour than fully parametric GLMs have been used, such as generalized additive models (GAM; Schmidt et al., 2010) that use non-parametric smooth functions to allow more flexibility in the relationship of response variable and predictors (Hastie et al., 2009). Machine learning approaches have also been successfully applied to wind disturbance modeling (see Hanewinkel et al. 2004 for an early example) and recently especially tree-based ensemble models, such as random forests, have been shown to perform well in predicting wind damage (Albrecht et al., 2019; Hart et al., 2019; Kabir et al., 2018; Schindler et al., 2016). While machine learning methods and additive models are able to more flexibly fit the data and account for non-linearities, the GLMs have strengths in their straightforward interpretability and the robustness of predictions (Albrecht et al., 2019; Nakou et al., 2016).

In this study, our goal was to create high-resolution spatial information about forest vulnerability to wind damage in Finland, using an extensive damage observation data set and a large compilation of spatial data sources to achieve this. More specifically, we aimed to (1) create a damage probability statistical model based on a large data set of wind damage observations in the Finnish National Forest Inventory (NFI), (2) compare three statistical and machine learning methods for creating the model: GLM, GAM and BRT, (3) calculate a damage probability map by combining the model with national extent GIS layers of model predictors, compiled from different sources, and (4) test the performance of the map with independent damage observations from new NFI data.

## 2. Material and methods

### 2.1 National Forest Inventory and wind damage observations

In this study, we used stand level wind damage observations from the 11^th^ Finnish national forest inventory (NFI11) to create an empirical model of wind damage probability (Fig. 1). The field work for the NFI11 was conducted from 2009 to 2013 (Korhonen, 2016; Korhonen et al., 2017). In later stages of the study, we also used NFI12 (field work in 2014 to 2018) to test the created map (see section 2.5).

**Figure 1.**
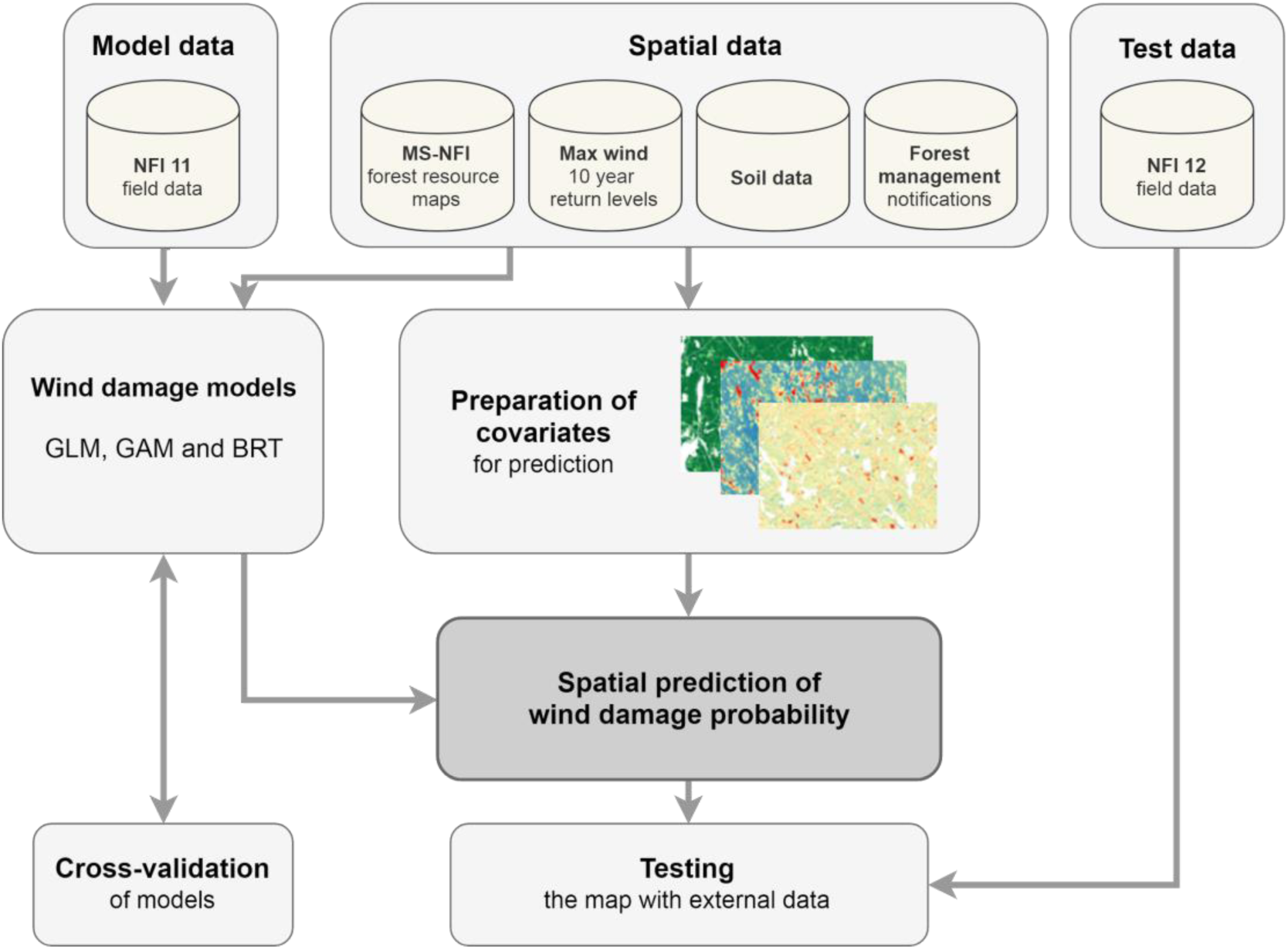
General approach and workflow

In our analysis, we only included plots that were defined as forest land. Poorly productive forests were excluded because they are unimportant for forestry and their wind damage risks tend to be small due to low volume of growing stock. In addition, plots on treeless stands or seedling stands without upper canopy layer were excluded because seedlings have very low wind damage probability (8633 plots). Plots with missing data or unrealistic (erroneous) values for any of the used variables were excluded (52 plots). Plots within less than 1 km from the national border were also excluded, as the data set describing local wind conditions (Venäläinen et al., 2017) had edge effects (214 plots). If a plot was located on the border of two or more forest stands, we only used the data from the stand where the plot centre was located. The final data set consisted of a total of 41 392 NFI plots.

Observations of stand level wind damage and an estimate of the damage time is documented in the Finnish NFI (Korhonen, 2016; Tomppo et al., 2011). Here, we used only the wind damage observations that had occurred no more than 5 years before the date of the field visit. Since the field work of NFI11 was done in 2009 to 2013, the data can contain observations from damage that has occurred between 2004 and 2013. During these years, several high impact storms affected Finland, such as cyclone Dagmar (known as Tapani in Finland) in December 2011 and a series of severe thunderstorms in summer 2010.

The severity of damage was not considered in the analysis, because the degree of damage was only recorded as cumulative effect of all damage agents, and no information of wind damage severity was available in cases where there were more than one damaging agent present. The restriction of the analysis to only severe damage cases would also have limited the number of damage observations available. Therefore, the binary damage variable contains stands with different damage severities. Stand level wind damage was observed at 1 070 plots of the total 41 392 NFI plots in the dataset.

### 2.2 Model predictors

#### 2.2.1 National Forest Inventory data

Most predictors in the statistical models were extracted from the NFI field data (Table 1 and 2). To describe the forest characteristics of the stand, dominant tree species and mean tree height in the stand were used. If several canopy layers and species were recorded in the data, the values from the layer with largest tree height were used, as the tallest trees can be assumed to be most vulnerable to wind. The NFI also documents the type and time of most recent forest management operation, and based on this data we created a variable describing the time since last thinning.

**Table 1.** Description of predictors used and their sources in the model and in the damage probability map. See section 2.2.1 for details.

| Variable | Type* | Unit / Classes | Source in model | Source in map |
| --- | --- | --- | --- | --- |
| Tree species | C | pine, spruce, other | NFI11 | MS-NFI 2015 |
| Tree height | N | dm | NFI11 | MS-NFI 2015 |
| Time since thinning | C | 0-5, 6-10, > 10 years | NFI11 | MS-NFI 2015, Forest use notifications |
| Wind (10-year return level of max wind speed) | N | ms<sup>-1</sup> | Venäläinen et al. 2017 | Venäläinen et al. 2017 |
| Open neighbor stand | C | True, False | MS-NFI 2013 | MS-NFI 2015 |
| Soil type | C | Mineral/coarse, Mineral/fine, Organic | NFI11 | GTK 2018, NLS 2018 |
| Mineral soil depth < 30 cm | C | True, False | NFI11 | GTK 2018, NLS 2018 |
| Site fertility | C | Fertile, Poor | NFI11 | MS-NFI 2015 |
| Temperature sum (average 1985-2014) | N | 100 dd (over 5C) | Aalto et al. 2016 | Aalto et al. 2016 |
| Damage density ratio | C | 0-2, 2-3, <3 | NFI11 | In the calculation of the map, this variable was included as a weighted average of all classes, because it was included in the model only to account for spatial structures in storm severity. |

\* C – categorical, N – numerical (continuous)

**Table 2.** Descriptive statistics for the NFI11 data. Mean and standard deviation for non-damaged, damaged and all plots continuous variables, and percentages of each class for categorical variables. The definitions of the variables are in table 1.

|  | Non-damaged | Damaged | All |
| --- | --- | --- | --- |
| Number of plots | 40322 | 1070 | 41392 |
| Species |  |  |  |
| *Scots pine* | 63.4% | 59.1% | 63.3% |
| *Norway spruce* | 24.0% | 36.8% | 24.3% |
| *Other* | 12.6% | 4.1% | 12.4% |
| Tree height | 163.0 (50.5) | 195.2 (45.1) | 163.9 (50.6) |
| Time since thinning |  |  |  |
| 0-5 years | 13.4% | 26.0% | 13.7% |
| 6-10 years | 9.2% | 15.5% | 9.4% |
| > 10 years | 77.4% | 58.5% | 76.9% |
| Wind | 12.1 (2.0) | 12.5 (2.0) | 12.2 (2.0) |
| Open neighbor |  |  |  |
| *False* | 85.7% | 84.6% | 85.7% |
| *True* | 14.3% | 15.4% | 14.3% |
| Soil type |  |  |  |
| *Mineral, coarse* | 66.9% | 77.8% | 67.2% |
| *Mineral, fine* | 12.7% | 9.8% | 12.7% |
| *Organic* | 20.3% | 12.4% | 20.1% |
| Soil depth < 30 cm |  |  |  |
| *False* | 89.5% | 85.0% | 89.4% |
| *True* | 10.5% | 15.0% | 10.6% |
| Site fertility |  |  |  |
| *Poor* | 34.8% | 31.9% | 34.7% |
| *Fertile* | 65.2% | 68.1% | 65.3% |
| Temperature sum | 1185 (178.9) | 1262.6 (130.4) | 1187.0 (178.3) |

NFI information about soil type, soil depth and site fertility was also used (Table 1 and 2). Soil type variable differentiated between organic and mineral soils, as well as fine and coarse grained mineral soils. Fine mineral soils included clay and fine sands, whereas sands and coarser soils were classified as coarse mineral soils. Grain size was estimated on the field by NFI teams. Site fertility classes in the NFI are estimated in eight classes, but in our analysis they were regrouped into two classes so that class “Fertile” contained sites from herb-rich to mesic forests on mineral soils and from euthrophic to meso-oligothrophic peatlands. Less fertile classes were included in the “Poor” fertility class (see Tomppo et al., 2011 for detailed description of the site fertility classes used in Finnish NFI).

The used data covers the whole country and contains damage observations from several years and several storm events. Therefore, not all plots were exposed to similar wind conditions and this needed to be taken into account in the statistical model. However, we did not have reliable data available about the spatial variation in maximum wind speed conditions during the study period and lacking such an important factor affecting the damage probability is likely to bias the estimation of the effects of other predictors. Therefore, a different approach was taken. To account for areas subjected to severe storm events, variable “Damage density ratio” was calculated using the locations of NFI plots as as the ratio of 2D kernel density of damaged plots and all plots (Table 1). That is, the ratio describes the spatial density of damaged plots in comparison to all NFI plots included in the model and a value of 2, for example, can therefore be interpreted as two times higher density of damaged plots than what would be expected from the density of all plots. The damage density variable was then transformed into a categorical variable (with classes 0-2, 2-3, and >3). The upper limit of the lowest class was set relatively high to identify only the strongest clusters of damaged plots and to avoid catching all the large-scale spatial trends with this variable. The calculations were done in R with the *KernSmooth* package (Wand, 2015) using bandwidth of 20 km, see details in S1.

#### 2.2.2 Other data sets and the delineation of forest stands

In addition to the NFI field data we also supplemented the model predictor set with additional variables describing local wind conditions and open forest borders from other data sources (Table 1 and 2). For the wind conditions, we used a data set describing the local 10-year return levels of maximum wind speeds in 20 × 20 m^2^ raster cells (Venäläinen et al., 2017). That is, the value of each pixel represents the level of maximum wind speed (ms^−1^) expected to be reached on average once in every 10 years. The data is downscaled from coarse-scale wind speed estimates in ERA-Interim reanalyzed data with a wind multiplier approach using CORINE land-use data and digital elevation model (Venäläinen et al., 2017). The data set contains maximum wind speeds calculated for eight different wind directions, and in this study we used the maximum value of these for each pixel. To identify stands with open forest borders (variable ‘Open neighbour stand’, Table 1), we used the multi-source NFI forest resource maps (MS-NFI; Mäkisara et al., 2016; Tomppo et al., 2008) that combine satellite data and NFI field data to create national extent forest resource maps in a 16 × 16 m^2^ resolution grid.

However, the used wind damage observations were documented on the level of forest stands and the stand borders were not mapped in the data but only estimated by the NFI team at the field. Therefore, in order to combine the stand-level damage information with other data sources, the locations of stand borders first needed to be defined. A forest stand in the the Finnish NFI is defined as spatially continuous land area that is homogeneous with respect to properties such as administrative boundaries, site fertility, structure of the growing stock (e.g. maturity class, tree species composition) and forest management (Tomppo et al., 2011). To create polygons that would approximately correspond to the stands assessed in the field by the NFI team, we used image segmentation on the MS-NFI data layers (corresponding to year 2013) describing growing stock volumes by main tree species groups (pine, spruce and deciduous species) and tree height. Land property boundaries obtained from the National Land Survey of Finland were also included in the segmentation, as they are considered as stand boundaries in the NFI. The image segmentation was conducted with the methodology described by Pekkarinen (2002), using the “segmentation by directed trees” algorithm by Narendra and Goldberg (1980).

Once the stand polygons were defined with image segmentation, they were used for calculating local wind conditions and finding stands with open stand borders. For each stand polygon, maximum wind-speed within the stand boundaries was calculated (Table 1). Maximum value was used because the NFI field data does not specify the exact location of the damage within the stand, and we assumed that damage occurred in the most wind exposed part of the stand.

To identify plots with open neighbor stands, median tree height was first calculated for each stand polygon using the MS-NFI tree height data. A stand was defined to have an open stand neighbor if the median tree height of any of the stand neighbours was smaller than 5 meters (Table 1). Median was used instead of mean so that it would be less affected by possible outlier values resulting from inaccuracies in defining the stand polygons.

Calculations of maximum wind speeds and open stand neighbors for the segments were conducted with PostGIS (version 2.4.0) and Python (version 2.7.12) with packages *geopandas* (version 0.3.0) and *rasterstats* (version 0.12.0).

### 2.3 Statistical modelling

Damage probability models were created using three different methods: generalized linear models (GLM), generalized additive models (GAM, Wood 2006) and boosted regression trees (BRT; Elith et al., 2008). In all the models the dependent variable was the presence of wind damage in the stand and independent variables described forest characteristics, forest management history, soil and site type, the 10-year return level of maximum wind speed and temperature sum (Table 1).

Binomial GLM with logit-link function were fitted in R (version 3.5.1, R Core Team, 2017). To account for non-linear relationships, logarithm transformation were tested for all continuous independent variables and included in the final model if they showed lower AIC than models with non-transformed variables. The transformations were included only for the GLM model, since GAM and BRT enable more flexibility in the shapes of the relationship between response variable and predictors, and can therefore account for non-linear relationships without transformations.

Variable selection was based on several criteria: (1) only variables that, based on earlier research, were expected to have a causal effect to wind damage probability were included, (2) since the ultimate goal of the model was to produce the damage probability map, we only included variables for which reasonably high-quality national-extent GIS data sets were available or could be derived from existing data, (3) the behaviour of the variable in the model was plausible based on existing understanding of forest wind damage. We also aimed to build the model so that all major components related to wind damage probability were included. Collinearity of predictors was inspected with Pearson’s correlation coefficients and generalized variance inflation factors (GVIF, Fox and Monette, 1992). All correlation between included continuous predictor variables were weaker than 0.5 and GVIFs for all variables were lower than 4.

Generalized additive model (GAM) is a generalized linear model with a linear predictor involving a sum of smooth functions of covariates. This specification of the model in terms of smooth functions instead of detailed parametric relationships allows for more flexibility in the dependence of the response of the covariates (Wood, 2017). In our analysis, GAM with logit-link function was fitted in R with package *mgcv* (version 1.8-24, Wood, 2011), using the same predictors that were included in the GLM. All continuous predictors were included in the model through non-linear smoothing spline functions. The dimension parameter (k), effectively setting the upper limit on the degrees of freedom related to the smooth, was set to 15 for all variables, except for temperature sum for which k=5 was chosen to avoid unrealistically fluctuating large-scale patterns in the predictions. The effective degrees of freedom (edf) after fitting the model were lower than k for all of the terms (see S2 for details), suggesting that the chosen k’s were sufficiently large.

Boosted regression trees (BRT) is an ensemble method, that combines a large number of regression trees with a boosting algorithm (Elith et al., 2008). Here, BRTs were computed with R package *dismo* (version 1.1-4, Hijmans et al., 2017). To find the best parameters, BRTs with different parameter combinations of tree complexity (tested values 1, 2, 3 and 5), learning rate (0.05, 0.01 and 0.005) and bag fraction (0.5, 0.6 and 0.75) were fitted. The number of trees was not assigned manually, but was estimated with k-fold cross-validation using the function *gbm.step* (Hijmans et al., 2017). To estimate the number of trees and to compare different parameter combinations, *gbm.step* was run separately for each parameter combination. Following the rule-of-thumb suggested by Elith et al. (2008), we excluded parameter combinations that led to models with fewer than 1000 trees. Thus, the model with parameter combination leading to lowest holdout residual deviance in the cross-validation performed by *gbm.step* and at least 1000 trees was chosen for the final model (tree complexity=2, learning rate=0.01, bag fraction = 0.5, 2250 trees, see Supplementary material for details).

To make sure that the unbalanced ratio of damaged versus non-damaged plots did not affect the results, BRTs were fitted also from two balanced datasets where the balancing of the observations was done by (1) undersampling the non-damaged plots or (2) oversampling the damaged plots. In both cases the cross-validated AUCs were very similar to ones calculated from the original unbalanced dataset and, therefore, the original data set was used for the final results.

To account for the sampling design, weights based on the forest area each plot represents were used in all models (Korhonen, 2016). For example, in northern Finland the NFI sampling design is sparser and therefore the weight of one plot in modelling is higher. To test if the clustered sampling design had an effect on the results, GLMs and GAMs were also fitted as mixed models (GLMM and GAMM) with plot clusters as random intercepts, using R packages lme4 (Bates et al., 2015) for GLMM and gamm4 (Wood and Scheipl, 2017) for GAMM. However, as the mixed model predictions (in the scale of the linear predictor, using only fixed effects for prediction) were highly correlated with the fixed effect model prediction (Pearson’s r=0.998, p<0.001 for GLM vs GLMM, and r=0.979, p<0.001 for GAM vs GAMM) and our interest was in marginal instead of conditional inference, no random effects were included in the final models.

The models were validated with 10-fold stratified cross-validation, where number of damaged plots was divided evenly into the folds. In the cross-validation, the variation in damage density variable was not used in the prediction, because the variable was included in the model only to account for spatial structures in storm severity in the data, and in an aimed use case of the models (i.e., estimating damage vulnerability in future events) we would not have this information available. Instead, separate predictions for test-folds were calculated with each class of the damage density variable (0-2, 2-3, >3). Then, these three predictions were averaged based on the frequency of each class in the original model data. See details in S1.

ROC curves and AUC values were calculated for each iteration of cross-validation and used to assess the performance of the models (see Supplementary material). The ROC curve plots the true positive rate (sensitivity) and true negative rate (specificity) of the model with all possible classification thresholds. The AUC values represent the area under ROC curve and measure the model’s ability to discriminate between events and non-events. AUC values of 0.5 corresponds to a situation where the classifier is no better than random (ROC curve along diagonal) and value of 1 a situation where the model perfectly discriminates between events and non-events. As a rule of thumb, AUC values over 0.7 are considered acceptable discrimination between classes, values over 0.8 excellent and values over 0.9 outstanding (Hosmer et al., 2013).

### 2.4 Calculation of the damage probability map

A GIS raster data layer with resolution of 16 × 16 m^2^ and extent of the whole country was prepared for each predictor variable used in the models (Table 1). Forest variables (dominant species, tree height, height-diameter ratio, open forest edge) were derived from the most recent Finnish MS-NFI data for year 2015 (Mäkisara et al., 2019). A grid cell was defined to be on an open forest edge if tree height in the MS-NFI data was lower than 5 meters in any of the cell’s within a 5 × 5 cell neighborhood.

Spatial data on forest management history (the time of last thinning) was derived from the forest use notification collected by the Finnish Forest Centre. This data consists of forest use notifications that forest owners are required to report to the Forest Centre before conducting management operations in their forests. For each 16 × 16 m^2^ pixel, we first assigned the year of the latest notification of planned thinning in that location of the pixel and then calculated the difference to year 2015.

Data for the 10-year return rates of maximum wind (Venäläinen et al., 2017) was resampled to the 16 × 16 m^2^ grid with GDAL using bilinear interpolation. Soil type was defined as ORGANIC for areas within the peatland polygons in the Topographic Database produced by the National Land Survey of Finland (NLS, 2018). Other areas were defined as mineral soils, and further divided to fine or coarse mineral soils based on the top soil information in the 1:200 000 resolution soil map of the Geological Survey of Finland (GTK, 2018). Data layer for soil fertility classes was made by reclassifying the MS-NFI fertility class data layer from the original five classes to the two classes used in the models (see details in section 2.2.1). Average annual temperature sum was calculated with a threshold of 5°C from daily weather data grids (Aalto et al., 2016) for the years 1985 to 2014.

Similarly as in the cross-validation, the variation in damage density variable was not used in the prediction, because we would not have this information available for future events. Instead, separate predictions were calculated with each class of the damage density variable and these three predictions were then averaged based on the frequency of each class in the original model data. See details in S1.

The damage probability map was calculated from the GLM, GAM and BRT model objects and the GIS data layers using R packages *raster* (Hijmans, 2017) and *sp* (Pebesma and Bivand, 2005).

### 2.5 Testing the map with new damage observations

The accuracy of the damage probability map was validated with an independent test data set. The map was compared to the damage observations in the most recent NFI measurements (12^th^ Finnish NFI, NFI12), which were not included in the model fitting data that was from the NFI11. Compared to NFI11, which covers the whole country, NFI12 does not cover the northernmost parts of Finland as plots in the three most northern municipalities (Northern Lapland), where the proportion of forest land is low, are not measured as frequently as other parts of the country.

We included the NFI12 plots that had been measured during 2014-2018, were classified as forest land by the field team, and were located within forest area in the MS-NFI forest resource maps (i.e., there were data in the wind damage probability map at the location of the plot). For wind damage we also used the same criteria as with the model data, i.e. only observations estimated to have occurred during the last 5 years were included and the severity of the damage was not considered. In addition, those permanent plots that were measured already in NFI11 were excluded from the test data, as the previous measurements in the same plots were used in the model fitting. The final test data consisted of 33 754 plots with wind damage in 734 of the plots.

Values of the wind damage probability maps were extracted at the locations of test data plots as the mean value of map pixels within 20 meter buffer from the location of the plot center. ROC curves and AUC values were calculated from the wind damage information in the test data and the extracted values of the damage probability maps. The extraction was conducted in R with package *raster* (version 2.8-19, Hijmans, 2017) and ROC/AUC calculations with package *pROC* (version 1.12.1, Robin et al., 2011).

## 3. Results

The results showed that forest vulnerability to wind damage is strongly driven by forest characteristics, especially tree height (Figs 2-4, Table 3). In all models, the damage probability increased with tree height, and the increase was strongest for spruce dominated forests. Also forest management affected damage probability in the models, as recently thinned forests and forests with open stand borders were more susceptible to damage. These predictors, related to the forest characteristics, very much drive the fine-scale spatial variation of damage probability in the (Fig. 7).

**Figure 2.**
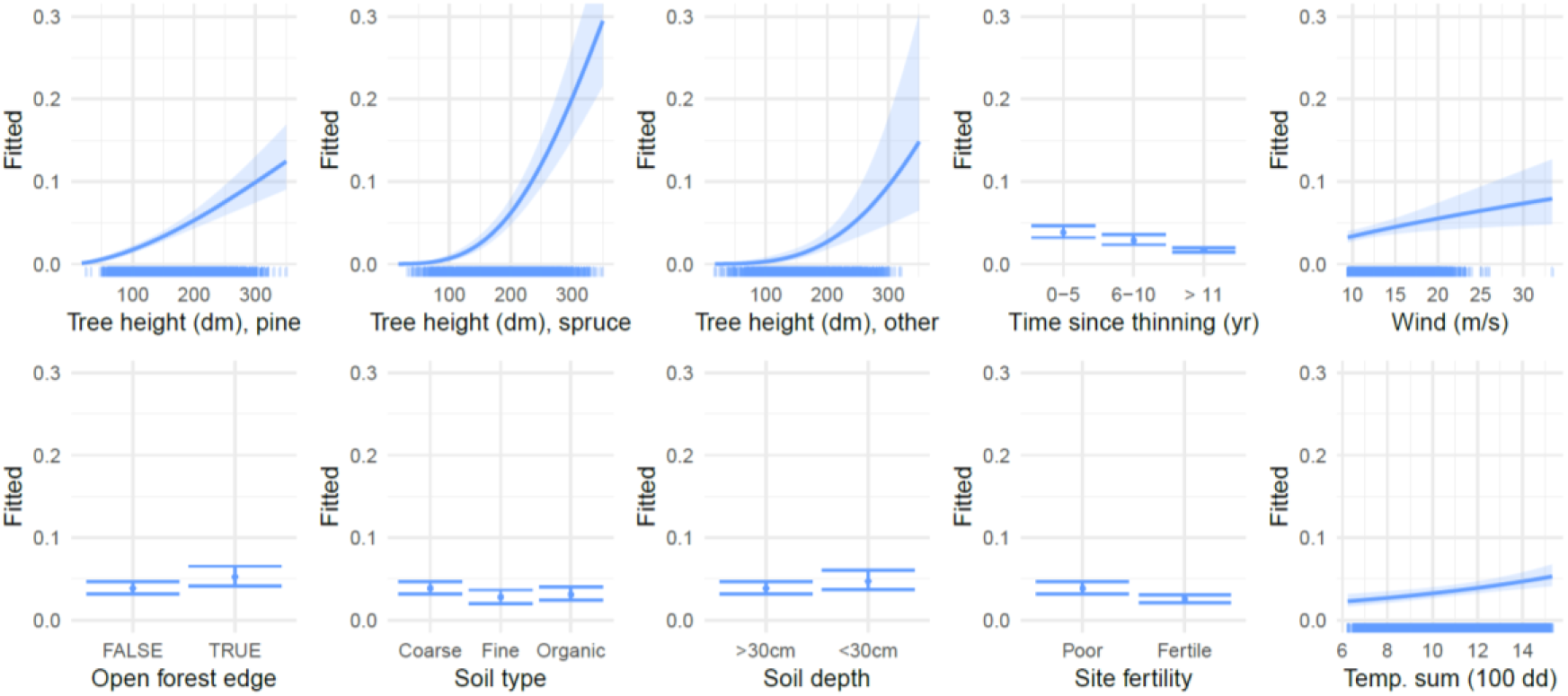
GLM partial dependence plots for the map predictors. Prediction of damage probability is calculated for the range of each predictor variable when other predictors are set to average (continuous variables) or reference class (categorical variables). Rugged x-axis describes the distribution of data. Confidence intervals are calculated as 2 x prediction standard error (in the scale of the linear predictor).

**Figure 3.**
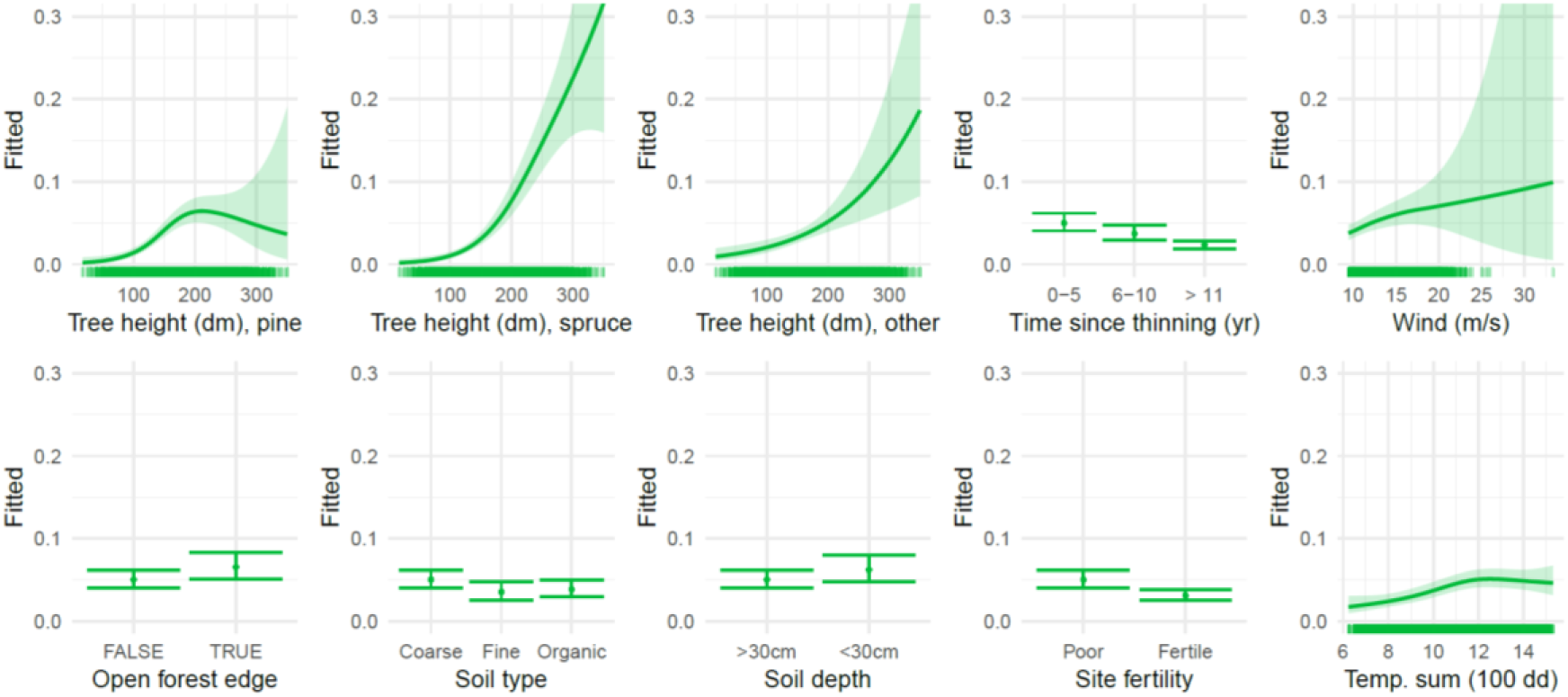
GAM partial dependence plots for the map predictors. Prediction of damage probability is calculated for the range of each predictor variable when other predictors are set to average (continuous variables) or reference class (categorical variables). Rugged x-axis describes the distribution of data. Confidence intervals are calculated as 2 x prediction standard error (in the scale of the linear predictor).

**Figure 4.**
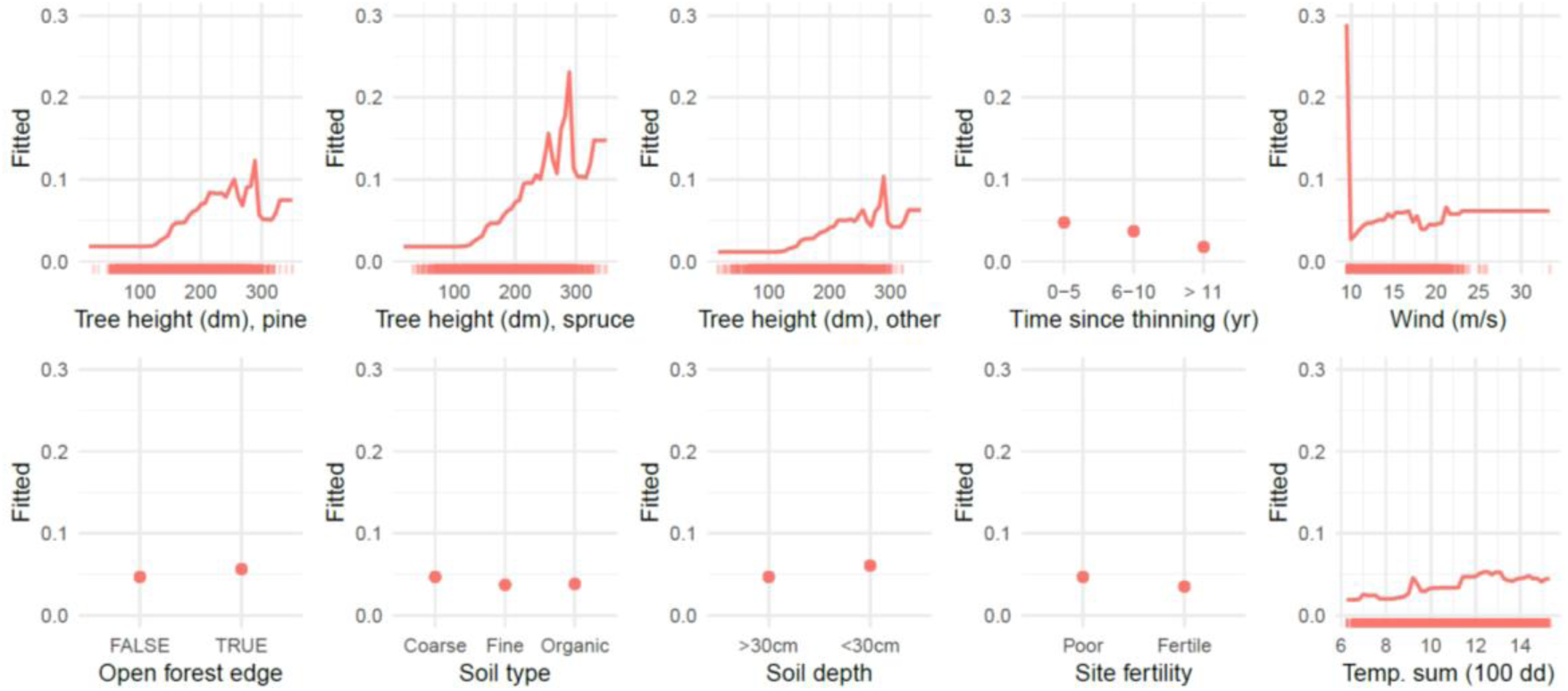
BRT partial dependence plots for the map predictors. Prediction of damage probability is calculated for the range of each predictor variable when other predictors are set to average (continuous variables) or reference class (categorical variables). Rugged x-axis describes the distribution of data.

**Table 3.** GLM model results (for categorical variables, the first class listed in Table 1 is the reference class, and therefore not listed separately in this table).

|  | Estimate | Std. Error | z value | Pr(>\|z\|) |
| --- | --- | --- | --- | --- |
| (Intercept) | -14.690 | 1.061 | -13.841 | < 0.001 |
| Species/Spruce | -8.494 | 1.918 | -4.430 | < 0.001 |
| Species/Other | -9.314 | 3.931 | -2.370 | 0.018 |
| log(Height) | 1.661 | 0.189 | 8.807 | < 0.001 |
| Last thinning/6-10 years | -0.298 | 0.113 | -2.637 | 0.008 |
| Last thinning/over 10 years | -0.844 | 0.084 | -9.995 | < 0.001 |
| log(Wind) | 0.749 | 0.238 | 3.152 | 0.002 |
| Open stand border / TRUE | 0.310 | 0.095 | 3.284 | 0.001 |
| Soil/mineral, fine | -0.356 | 0.124 | -2.875 | 0.004 |
| Soil/organic | -0.216 | 0.110 | -1.962 | 0.050 |
| Soil depth < 30cm / TRUE | 0.214 | 0.106 | 2.011 | 0.044 |
| Site fertility / Fertile | -0.425 | 0.092 | -4.611 | < 0.001 |
| Temperature sum | 0.096 | 0.025 | 3.843 | < 0.001 |
| Damage density / 2-3 | 1.104 | 0.088 | 12.498 | < 0.001 |
| Damage density / >3 | 1.898 | 0.111 | 17.137 | < 0.001 |
| Species/Spruce : log(Height) | 1.634 | 0.358 | 4.561 | < 0.001 |
| Species/Other : log(Height) | 1.625 | 0.742 | 2.190 | 0.029 |

Wind damage probability was found to show distinct large-scale trends, most importantly the decreasing damage probability from south to north (Fig. 7). This effect in the models comes from the temperature sum, but also other predictors contributed to the large-scale trends in the map, as there as large-scale patterns in wind conditions, forest characteristics and soil and site fertility conditions (Figs 2-4). The north-south pattern in damage density was evident in the damage probability maps with all model methods. However, the map created with the BRT model showed unexpectedly high damage probability values for the northernmost parts of the country (Fig. 7).

The model predictors showed in general rather similar effects in the three tested methods (GLM, GAM and BRT). Yet, there are also differences, especially in the shape of relationship between the continuous predictors and predicted damage probability (Figs 2-4). In GLM, the relationships are restricted to sigmoidal curves, whereas GAM and BRT allow more flexible shapes of responses. This can be seen, for example, in how increasing tree height in pine forests shows steadily increasing damage probability with GLM (Fig. 2) whereas in GAM damage probability peaks around tree height 200 dm and then declines. Higher values of damage density ratio led to higher damage probability in all models, as expected (Fig. 5).

**Figure 5.**
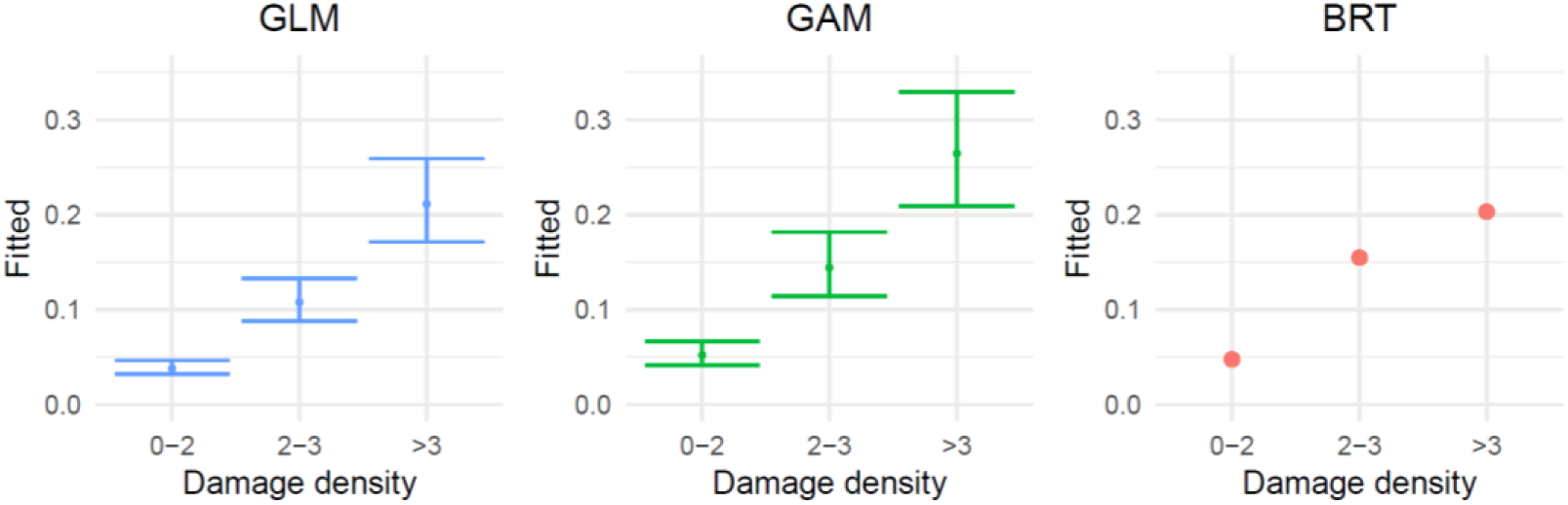
Partial dependence plots for damage density in the different models (GLM, GAM and BRT). Damage density was included in the models to account for spatial variation in severity of storm damage in the data, and it was set to 0 when calculating the wind damage probability map. Note that the y-axis range differs from figures 2-4.

As the BRT predictions are calculated from ensembles of regression trees, they enable very sharp changes in the prediction within small changes in the predictor (Fig. 4). They can also contain diverse interactions between the predictors, which are unfortunately not visible in partial dependence plots like Fig. 4. The BRT results showed somewhat different trends than the other methods in model responses to predictors (Fig. 4). For example, while tree height in spruce forests increases damage probability throughout the range of data in GLM and GAM results (Figs 2-3), in BRT results similar strongly increasing trend is not found, instead the relationship between height and damage probability seems to saturate for all tree species (Fig. 4). The large-scale spatial patterns in map prediction also differed for BRT compared to the other models, as high values of damage probability were predicted for the northernmost parts of the country. (Fig. 7).

Cross-validation showed higher predictive performance of the GAM model compared to the GLM and BRT (Fig. 6). However, when the final damage probability maps were tested with the NFI12 test data, all models showed very similar performance in discriminating between damaged and non-damaged plots in the test data. (Fig. 8). All maps gave on average higher damage probability values for damaged than non-damaged plots and showed an acceptable level of discrimination between the two (AUC > 0.7). The added flexibility and ability to account for nonlinear relationships in GAM and BRT did not considerably improve the predictive performance of maps compared to the fully parametric GLM (Fig. 8).

**Figure 6.**
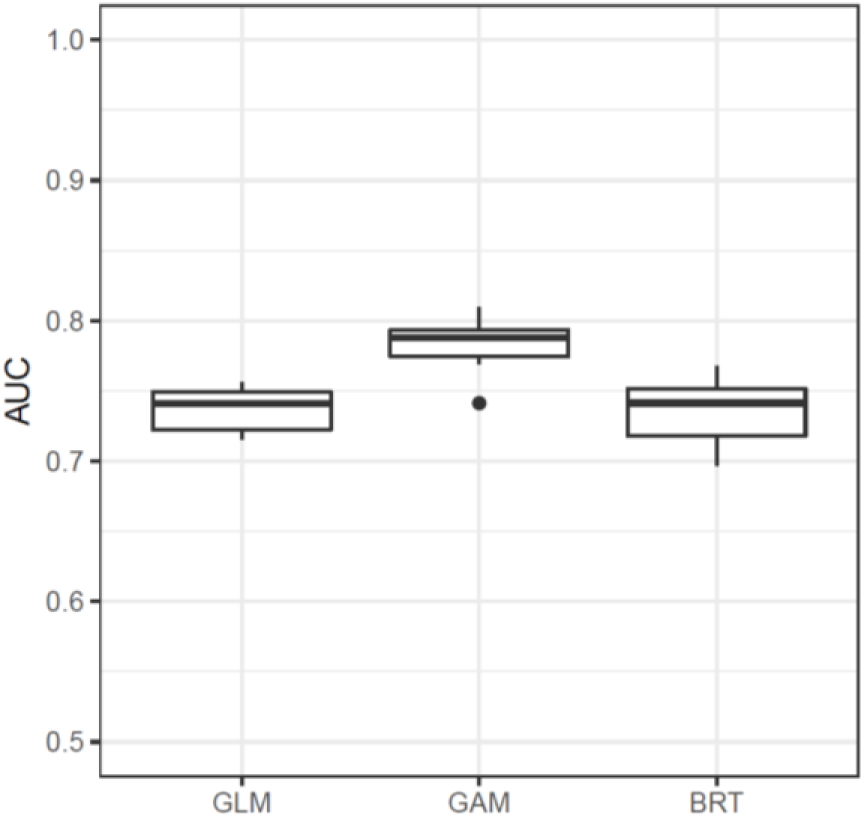
Distribution of AUC values in the 10-fold cross-validation for GLM, GAM and BRT.

**Figure 7.**
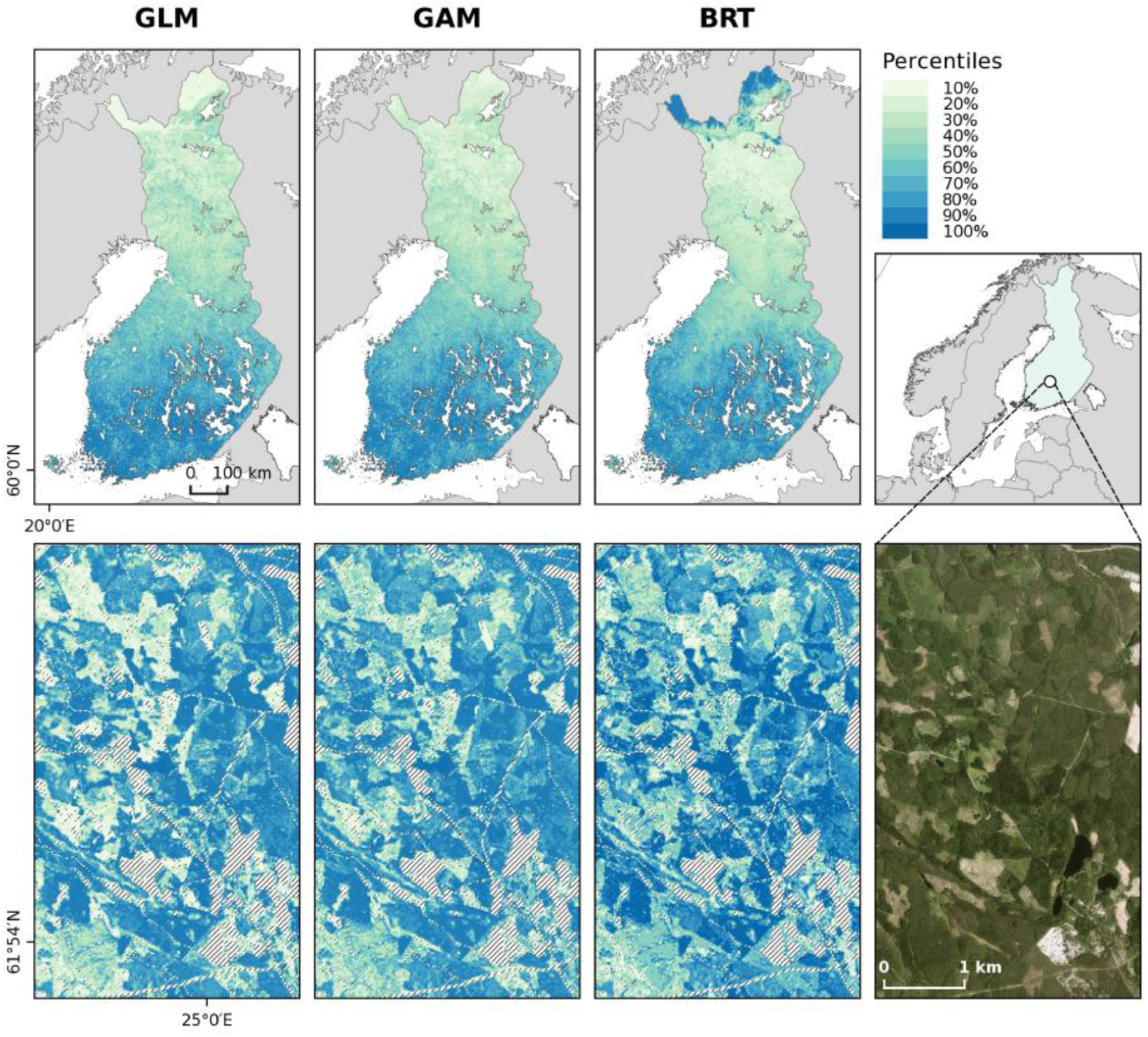
Damage vulnerability maps calculated for the whole country (upper panel) and a fine-scale detail of the maps (lower panel), calculated with the three different damage probability models (GLM, GAM and BRT), and an orthophoto from the same location (B). Colors in the damage vulnerability map are defined by the percentiles of the map data (e.g., the first class contain the lowest 10% of map values). The upper panel maps are resampled to 1 km × 1 km resolution with bilinear interpolation. Note that the orthophoto is not from the exact same time as the forest resource data used for the calculation of the map. Orthophoto © National Land Survey of Finland.

**Figure 8.**
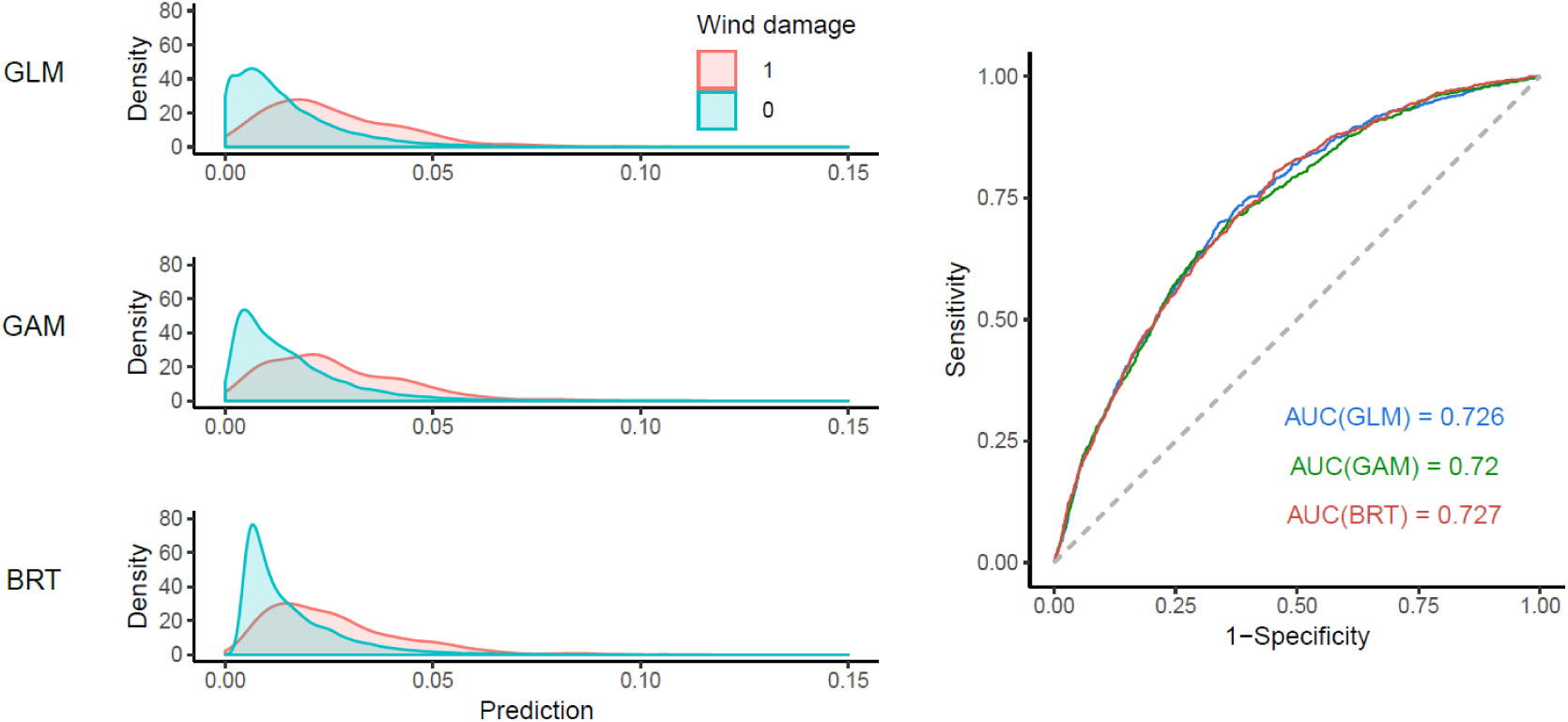
Density plots of the distributions of map predictions for test data plots with wind damage (red) and without wind damage (blue), and ROC curve showing the ability of the maps to distinguish between damaged and non-damaged test plots for the different model methods (GLM, GAM and BRT).

## 4. Discussion

### 4.1 The damage probability map

We created a new spatial wind damage risk product based on inventory data spanning over several years and several other data spatial sources, including information where the actual harvests have recently occurred in Finland. Validation of the map with independent and large data dataset showed that the map is able to identify vulnerable stands also in new storm events. While there have been attempts to map wind damage probability based on empirical damage models (Saarinen et al., 2016; Schindler et al., 2009; Suvanto et al., 2016), our work here uniquely provides national extent and high spatial resolution information about forest vulnerability to wind and is also tested with large external test data.

The successful identification of damage vulnerability in the independent test data is not trivial. First of all, wind damage is challenging to predict and extending the performance of statistical wind damage models to new data sets has been shown not to be straightforward (Fridman and Valinger, 1998; Kamimura et al., 2015; Lanquaye-Opoku and Mitchell, 2005). Moreover, because we wanted to test how well our map identifies forest vulnerability to wind in future events, for which we don’t have detailed information of, we did not include any information about spatial distribution of wind speeds or storm events during the time frame of the test data when we tested the map. Thus, the discrimination of damaged from non-damaged plots with fair accuracy (AUC=0.72) for the entire extent of Finland indicates that the map is indeed successful in identifying the vulnerable forests, and implies that efficient combination of inventory data and several new spatial data sources is a promising way to map damage risks.

A major factor contributing to the successful extension of the map to new test data was the large and systematically sampled forest and damage data that spanned over several years. Thus, our model was able to represent the different conditions (forest characteristics, soil, etc.) within the country. The need for comprehensive model data in empirical wind damage models has been demonstrated, for example, by Hart et al. (2019) who showed that it is possible to generalize to new storm events when the model data covers the variation of predictor variables in the new data set.

In addition to good representation of environmental and forest conditions, our data also represents different types of wind events, since the data consisted of damage observations in a 5-year time window. Most wind disturbance studies typically concentrate on one or few storms (e.g., Hart et al., 2019; Kamimura et al., 2015; Saarinen et al., 2016; Schindler et al., 2009; Suvanto et al., 2016), which limits their ability to generalize to different storm events. While modelling of multi-event data can be more challenging than single-event data (Albrecht et al., 2019), we argue that it is necessary when the purpose of the model is in assessing damage probability in future events.

Availability of high-quality and high-resolution spatial data of the model predictors was also crucial in the successful creation of the damage probability map. Additional uncertainties arise from the input data sets when model predictions are made with GIS data gathered from several different sources instead of the field-measured data that were used for fitting the model. In our case, we were able to utilize several high-quality and high-resolution data sources, such as the MS-NFI raster maps of forest characteristics (Mäkisara et al., 2019) and new data products of local wind conditions (Venäläinen et al., 2017). We were also able to use the recently opened forest use notification data from the Finnish Forest Centre that provided us with nation-wide information about the recent forest management history of the stands. This type of legacy information about forest management is typically difficult to obtain and has rarely been included in predictive wind damage risk models before, despite the clear effects of management history on forest disturbance dynamics. While all these data sources contain uncertainties, the verification of our map with independent test data showed that they were nevertheless able to represent well the main factors determining forest susceptibility to wind.

With new data sources and increasing quality and availability of data in the future, the accuracy of the map could still be improved. This could mean, for example, improved accuracy of tree height information through the use of lidar data or inclusion of variables that were left out of the current map due to lack of national level spatial data about their distribution (e.g. distribution of wood decaying fungi that weaken trees’ resistance to wind). Soil data had maybe the lowest resolution and higher uncertainties of the used GIS data and, therefore, increased quality of those data sets would also be desirable. However, the effects of soil variables in the model were relatively small, and therefore the effects of only improving the soil GIS data in the prediction would most likely not be drastic. Instead, more detailed soil data would be needed for the model data to improve the description of the role of soil characteristics on tree vulnerability to wind in the model.

### 4.2 Drivers of forest susceptibility to wind disturbance

The factors that were found to affect damage probability in our results are well in line with previously published results. For example, increasing damage probability with tree height and the higher vulnerability of Norway spruce have been shown in previous studies (Peltola et al., 1999; Suvanto et al., 2016; Valinger and Fridman, 2011). New stand edges after clearcutting of the neighboring stand and recently thinned stands have also been known to be at higher risk of windthrow (Lohmander and Helles, 1987; Peltola et al., 1999; Wallentin and Nilsson, 2014).

While open stand edges did increase the risk of wind damage in our results, the effect was not as distinct as could be expected from earlier research that emphasizes the role of forest edges (e.g., Peltola et al., 1999). This may in part result from the use of stand level data, where defining and identifying the open stand borders from the NFI data is more uncertain than in the case of tree-level analysis (see section 2.3.2 for the used methodology). Earlier work with storm damage data from severe autumn storms in Finland showed that the effects of open forest edges on damage probability were more emphasized in tree-level analysis (Suvanto et al., 2018) than in the stand-level analysis of the same data (Suvanto et al., 2016). In the future, potential improvements to the presentation of damage probability at the forest edges in the map could be achieved by combining tree-level results or mechanistic approaches to the current stand-level modeling approach.

In the model, the effect of wind speed data (Venäläinen et al., 2017) on damage probability showed logical behaviour of increasing damage probability with increasing 10-year return rates of maximum wind speed. The wind speed data accounts for the effects of topography on general wind conditions, and therefore variables describing topographical conditions were not included in our models, even though they have been shown to be linked with wind damage probability (e.g., Schindler et al., 2009).

Large-scale geographical patterns in our results showed that the probability of wind damage in Finland decreases from south to north. This is in agreement with results form previous studies combining forest model simulations with mechanistic wind damage models (Ikonen et al., 2017; Peltola et al., 2010). The higher susceptibility of forests in southern Finland to wind disturbances is related to the shorter length of the soil frost period in southern parts of the country. When the soil is frozen, trees are well anchored to the ground and less vulnerable to windthrow and, therefore, forests located in areas with longer periods of soil frost are less likely to be damaged during winter storms (Gregow et al., 2011; Laapas et al., 2019) (Gregow et al., 2011). However, other factors affecting forest wind susceptibility also change along the north-south gradient. The proportion of Scots pine, a species more resistant to wind than Norway spruce, increases towards north, and trees in the north have on average lower height-to-diameter ratio, which is linked to wind damage sensitivity (Ikonen et al., 2017; Peltola et al., 2010). In addition, in southern parts of the country, forest stands are smaller in area and there are less protected areas compared to the north. Thus, more frequent windthrows related to new stand edges and recent thinnings may also contribute to higher damage probability in the south. Similarly, butt rot caused by *Heterobasidion* sp., which increases tree vulnerability to wind (Honkaniemi et al., 2017), currently affects the southern parts of the country more severely (Mattila and Nuutinen, 2007; Müller et al., 2018) and may also contribute to the north-south pattern in the wind damage probability in our results. Therefore, it is not entirely clear what are the exact mechanisms causing increased damage probability with temperature sum in our model.

### 4.3 Comparison of methods

While the results for GLM and GAM models were rather similar, the BRT showed rather different model behaviour and large scale prediction patterns. The lack of test data in the northernmost parts of the country makes the interpretation of the test results (Fig. 8) for the BRT a bit challenging, as the area with unexpected BRT predictions is mainly not covered by the test data. In any case, the high values of BRT predictions in northernmost Finland do not seem realistic.

Our results did not show improved predictive performance of the map with the more flexible methods GAM and BRT compared to the logistic regression model (GLM). This is somewhat surprising, especially in the case of BRTs, because several recent studies have shown good performance of random forest for modelling storm disturbances (Albrecht et al., 2019; Hart et al., 2019; Kabir et al., 2018). Yet, in our results BRT did not lead to better predictive performance in cross-validation or with test data, even though it is a tree-based ensemble method very similar to random forest.

Our analysis differs from that of these earlier studies (Albrecht et al., 2019; Hart et al., 2019; Kabir et al., 2018) on a few aspects. First, we modelled wind damage on the level of forest stands, whereas the above mentioned studies were operating on tree-level. Second, we were using longer term NFI damage observations whereas most others used data from specific storm events. However, the study by Albrecht et al. (2019) contained both event-specific and non-event-specific data and they found random forests to outperform GLMs in both types of data. Third, we performed the cross-validation without considering the spatial variation in the storm conditions (the damage density variable in our analysis). This was done because we did not want to use this variable in the prediction, as the final aim was to generalize the results to future damage events, where this information would not be available. It is possible that this approach is disadvantageous to the BRT. All these differences in the approaches and analysis may have contributed in different performance of methods between the studies.

On the other hand, while the above mentioned studies did find machine learning methods outperform traditional statistical models in many ways, they also showed some positive sides of the logistic models. Most importantly, even though random forests showed superior performance when cross-validating models with data from one storm event in Hart et al. (2019), logistic models showed the highest AUC values compared to the other methods when the model was applied to another storm event, supporting the value of GLMs when generalizing the results to new storm events.

It seems that while machine learning methods such as BRT and random forest have advantages in accounting for more complex relationships and interactions in the data, they also catch patterns that are not helpful in estimating future disturbance probabilities (see, e.g., the unrealisticly high probabilities of damage with very low wind speeds in BRT, Fig. 4). This is likely to hamper the performance of BRTs so that they are not able to improve cross-validation performance compared to GLM.

Use of GLMs has the extra benefit of being more easily communicated to the end user, and they can be easily applied to new use cases when model coefficient estimates are published. The interpretation of relationships between predictors and the response variable is more straightforward, whereas especially in BRTs very small changes in e.g. tree height can lead to drastic changes in model prediction (Fig. 4). The unexpectedly high damage probability values in northern Finland also demonstrate the unpredictability of BRT model behaviour. This aspect is particularly important when the end product is meant to be used in practical applications.

### 4.4 Applications and use of the maps

The strength of the map is in its high resolution and large extent. The high-resolution makes it useful for assessing wind damage susceptibility of individual forest stands in fragmented forest landscapes where spatial variation of forest characteristics is high. On the other hand, the national extent of the map makes it widely available and accessible to everyone who is making forest management decisions in Finnish forests. To further improve the accessibility and usability of the map, we created an openly available web map application, where users can explore the map and find the estimated wind damage vulnerabilities of the forests they are interested in, without expert knowledge in GIS software (see https://metsainfo.luke.fi/en/tuulituhoriskikartta, currently only in Finnish, click “Tuulituhoriskit” box to see the wind damage vulnerability map). By providing an effective tool for identifying the vulnerable stands and for communicating wind damage risks to forest managers and owners, the map has potential to steer forest management practices towards a more disturbance-aware direction.

In addition to forest management, high-resolution information about forest wind vulnerability is crucially needed also in other sectors and applications. For example, the map can help in identifying high-risk locations where windthrown trees can harm infrastructure by damaging power lines and blocking roads. Insurance companies may also use high-resolution vulnerability information for a more risk based pricing of forest insurances.

While wind disturbances have major consequences from the human point of view, they are a natural process and have an important role in shaping the structure and function of forest ecosystems (Bouget and Duelli, 2004; Kuuluvainen, 2002). By exploring the drivers and spatial variability of wind disturbance dynamics, our results can therefore provide insight in current disturbance regime and its effects in the ecosystem, such as biodiversity and carbon cycling. Improved information about forest disturbances and tree mortality is also urgently needed for vegetation models from stand to global scales to understand how forests will react to the changing climate (Bugmann et al., 2019; Friend et al., 2014).

When applying the map in practice, it is important to consider its limitations. First, the damage probabilities in the map are in reference to the damage happened during the study period. The amount of wind damage varies strongly between years and future conditions are not likely to exactly match the conditions during the period from which the data comes from. Therefore, instead of exact probability values, it is better to interpret the map values as relative differences in damage vulnerability. Second, it is important to note that the damage probabilities do not only refer to complete damage of the stand, as our analysis also included less severe damage cases and we did not account for damage severity. Third, it is good to keep in mind that the map presents forest vulnerability to wind and it is not possible to predict the exact location of future wind disturbances, as there are many things - such as tracks and meteorological conditions of future storms - that can’t be accounted for in the map. The uncertainties need to be taken into consideration when using the map.

Wind disturbances are strongly linked to other processes of the forest and, therefore, should be considered in larger context. Thus, the greatest benefits of our results can perhaps be achieved by combining it with information and understanding of other processes that control forest ecosystems and forest management decisions. For example, the risk model can be coupled with forest growth simulators and thereafter storm damage risks of different forest management strategies can be evaluated simultaneously when making future scenarios of forests. The map can be combined with spatial information of wood volumes and prices to assess economic risks wind disturbances. Combining wind disturbance results with the dynamics of other disturbance agents is also crucial, as wind damage is strongly linked to bark beetle outbreaks and root rot, and these interactions are becoming increasingly important with the changing climate (Seidl et al., 2017; Seidl and Rammer, 2017). A comprehensive approach is therefore needed to understand and effectively manage wind disturbances in forests.

## 5. Conclusions

In this study, we show how probability models based on NFI damage observations combined with existing spatial datasets can be used to provide a fine-scale large-extent map of wind disturbance probability. We also demonstrate the ability of the map to identify vulnerable stands in future events with an extensive external test data. These maps provide a powerful tool for supporting disturbance-aware management decisions, communicating disturbance risks to forest owners, and accounting for the effects of windthrown trees in other sectors, such as maintenance of powerline infrastructures.

Our results show that machine learning methods, such as BRT, do not always provide superior results compared to traditional statistical models. As their interpretation in also less straightforward, they can sometimes lead to unpredictable prediction outcomes. Therefore, it is crucial to always assess the benefits of different approaches and to carefully test the performance of the used method with test data that is not used in model fitting. Partial dependence plots and other ways for exploration of model predictions in different situations also provide useful tools for assessing if model behaviour is realistic and biologically plausible.

The success of our results is based on large and representative model data as well as high-quality and high-resolution GIS data used as map inputs. In Finland, good data sets for both the model fitting and the map inputs are available, which enabled work done in this study. However, with improving data quality and availability (for both damage observations for model fitting and GIS data for map inputs), similar work could be extended to other regions and even to other disturbance types.

## Acknowledgements

The research was funded by Finnish Forest Foundation (project MyrskyPuu). We thank Ari Venäläinen and Mikko Laapas from the Finnish Meteorological Institute for their advice and the maximum wind speed 10-year return level data, and the MyrskyPuu project steering group (Liisa Mäkijärvi, Erno Järvinen, Heli Peltola, Ari Venäläinen and Eero Mikkola) for the discussions and their insights on the topic. We also thank Kari T. Korhonen for his comments on the manuscript and the whole NFI team in Luke for the NFI data we were able to use in the study. We acknowledge CSC – IT Center for Science, Finland, for computational resources.

## Supplementary materials

S1. The damage density ratio variable

S2. GAM model results

S3. BRT parameter tuning

S4. GLM variance-covariance matrix

