## Supplementary material S4 for "High-resolution mapping of forest vulnerability to wind for disturbance-aware forestry and climate change adaptation"

### Supplementary materials S1 – S3

#### S1. The damage density ratio variable

##### S1.1 Calculating the damage density variable

A. 2D kernel density of **all NFI11 plots included in the model fitting** was calculated with a bandwidth of 20 km.
B. 2D kernel density of the **plots with wind damage observations** was calculated with a bandwidth of 20 km
C. **Density damage ratio** was calculated as the ratio between the density of damaged plots and the density of all plots as *density(damaged plots) / density(all plots)*
D. The density damage ratio was reclassified to three categories: 0-2, 2-3 and > 3.

The calculations were carries out with R packages *KernSmooth* (Wand 2015) and *raster* (Hijmans 2017).

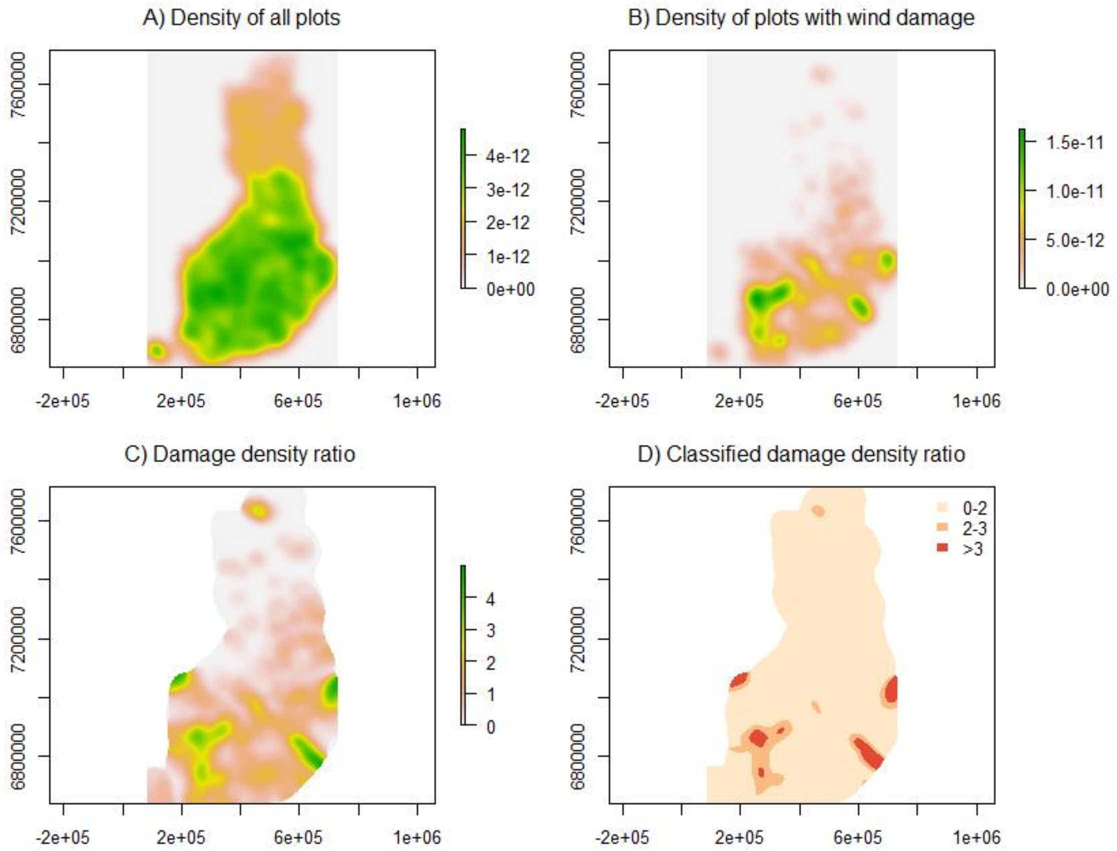


**Fig. S1.1.** Demonstration of phases in the calculation of damage density ratio.

#### S1.2 Predicting with the damage density variable

We did not want to include spatial variation in the damage density variable in the prediction (in cross-validation or in calculation of the maps) because it was only included in the model to account for spatial variation in storm frequency and conditions, and we would not have that information when trying to use the maps for assessing future vulnerability of forests to wind.

Therefore, when the model was used for prediction, predictions with all damage density classes (0-2, 2-3, >3) were performed separately for the whole data set and then averaged based on the frequency of each class in the original model data set:

$$P(damage)_{final} = \sum_{i=1}^{n} w_i \, P(damage)_{dd=i}$$

where:

| | |
|---|---|
| $P(damage)_{final}$ | is the final prediction for the probability of damage |
| $i$ | is the index for the classes of the damage density variable (1 = class ‘0-2’, 2 = class ‘2-3’, 3 = class ‘>3’) |
| $w_i$ | is the frequency of $i$th damage density class in the original data (0.905, 0.072, 0.023) |
| $P(damage)_{dd=i}$ | is the prediction of damage probability with $i$th class of damage density |

In calculation of the frequencies of the damage density classes in the original data ($w_i$) the NFI11 sampling design was taken into account by weighting each plot according to the forest area the plot represents (see Tomppo et al. 2011 for details).

#### S2. GAM model results

Family: binomial
Link function: logit

Formula:
wind_damage ~ s(Height, by = Species, k = 15) + Last_thinning + s(Wind_max, k = 15) + Open_stand_border + Soil + Soil_depth + Site_fertility + s(Temperature_sum, k = 5) + Damage_density

Parametric coefficients:

| | Estimate | Std. Error | z value | Pr(>\|z\|) |
|---|---|---|---|---|
| (Intercept) | -3.41015 | 0.10101 | -33.759 | < 2e-16 *** |
| Last_thinning_6_to_10y | -0.30609 | 0.11304 | -2.708 | 0.00677 ** |
| Last_thinning_over_10y | -0.81027 | 0.08539 | -9.489 | < 2e-16 *** |
| Open_stand_border_TRUE | 0.27623 | 0.09493 | 2.910 | 0.00361 ** |
| Soil_Mineral_fine | -0.38338 | 0.12317 | -3.112 | 0.00186 ** |
| Soil_Organic | -0.28075 | 0.10964 | -2.561 | 0.01045 * |
| Soil_depth_Under30cm | 0.22053 | 0.10671 | 2.067 | 0.03876 * |
| Site_fertility_High | -0.50363 | 0.08669 | -5.809 | 6.28e-09 *** |
| Damage_density_Class2-3 | 1.08956 | 0.08950 | 12.173 | < 2e-16 *** |
| Damage_density_Class>3 | 1.84798 | 0.11231 | 16.454 | < 2e-16 *** |

---
Signif. codes: 0 ‘***’ 0.001 ‘**’ 0.01 ‘*’ 0.05 ‘.’ 0.1 ‘ ’ 1

Approximate significance of smooth terms:

| | edf | Ref.df | Chi.sq | p-value |
|---|---|---|---|---|
| s(Height):Species_Pine | 2.888 | 3.677 | 93.51 | < 2e-16 *** |
| s(Height):Species_Spruce | 2.173 | 2.746 | 225.45 | < 2e-16 *** |
| s(Height):Species_Other | 1.001 | 1.003 | 15.69 | 7.56e-05 *** |
| s(Wind_max) | 1.934 | 2.469 | 16.21 | 0.00076 *** |
| s(Temperature_sum) | 2.562 | 3.098 | 24.39 | 2.40e-05 *** |

---
Signif. codes: 0 ‘***’ 0.001 ‘**’ 0.01 ‘*’ 0.05 ‘.’ 0.1 ‘ ’ 1

R-sq.(adj) = 0.0331 Deviance explained = 13.2%
UBRE = -0.81704 Scale est. = 1 n = 41392

### S3. BRT parameter tuning

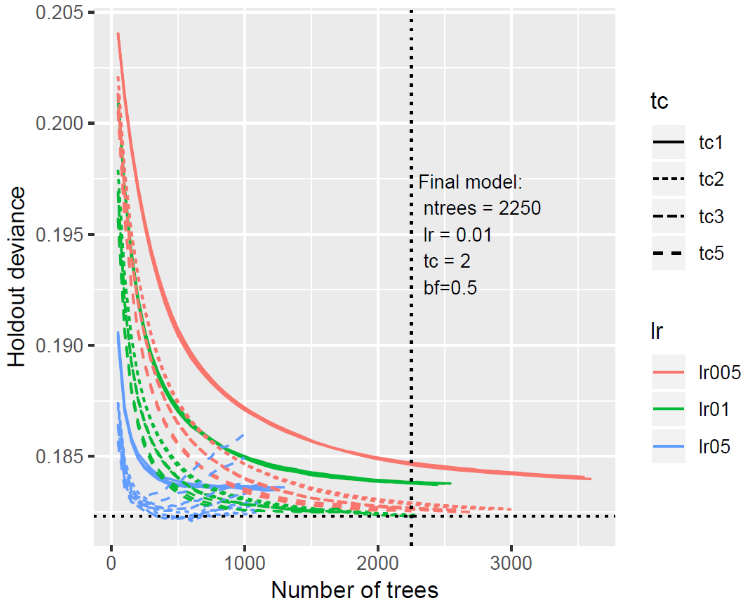


Grid search for different options of tree complexity (tc: 1, 2, 3 and 5), learning rate (lr: 0.005, 0.01, 0.5), and bag fraction (bf=0.5, 0.6 and 0.75) was performed. Different tr values are shown with different line types and lr values with different colors. Bag fractions are plotted for each combination of lr and tc but without distinct line type and color.

Number of trees fitted was chosen with *gbm.step* (*dismo* package, Hijmans et al. 2017). Holdout deviance describes the average residual deviance of the test data in cross-validation. Models with less than 1000 trees were excluded following the advice from Elith et al. (2008). Then, the parameter combination leading to smallest holdout residual deviance was chosen.

Black dotted lines show the final model location in the plot (number of trees = 2250, learning rate = 0.01, tree complexity = 2, bag fraction = 0.5).
